# Genomic consequences of colonisation, migration and genetic drift in barn owl insular populations of the eastern Mediterranean

**DOI:** 10.1101/2021.06.10.447833

**Authors:** Ana Paula Machado, Alexandros Topaloudis, Tristan Cumer, Eléonore Lavanchy, Vasileios Bontzorlos, Renato Ceccherelli, Motti Charter, Nikos Kassinis, Petros Lymberakis, Francesca Manzia, Anne-Lyse Ducrest, Mélanie Dupasquier, Nicolas Guex, Alexandre Roulin, Jérôme Goudet

**Affiliations:** Department of Ecology and Evolution, University of Lausanne, Lausanne, Switzerland; Green Fund, Kifisia, Athens, Greece; “TYTO” - Organization for the Management and Conservation of Biodiversity in Agricultural Ecosystems, Larisa, Greece; Centro Recupero Rapaci del Mugello, Firenze, Italy; Shamir Research Institute, University of Haifa, Katzrin, Israel; Department of Geography and Environmental Sciences, University of Haifa, Haifa, Israel; Game and Fauna Service, Ministry of the Interior, Nicosia, Cyprus; Natural History Museum of Crete, University of Crete, Herakleio, Greece; Centro di Recupero per la Fauna Selvatica–LIPU, Rome, Italy; Lausanne Genomic Technologies Facility, Lausanne, Switzerland; Bioinformatics Competence Centre, University of Lausanne, Lausanne, Switzerland; Swiss Institute of Bioinformatics, Lausanne, Switzerland

**Keywords:** Demographic inference, Inbreeding, Population genomics, *Tyto alba*, Whole genome sequencing

## Abstract

The study of insular populations was key in the development of evolutionary theory. The successful colonisation of an island depends on the geographic context, and specific characteristics of the organism and the island, but also on stochastic processes. As a result, apparently identical islands may harbour populations with contrasting histories. Here, we use whole genome sequences of 65 barn owls to investigate the patterns of inbreeding and genetic diversity of insular populations in the eastern Mediterranean Sea. We focus on Crete and Cyprus, islands with similar size, climate and distance to mainland, that provide natural replicates for a comparative analysis of the impacts of microevolutionary processes on isolated populations. We show that barn owl populations from each island have a separate origin, Crete being genetically more similar to other Greek islands and mainland Greece, and Cyprus more similar to the Levant. Further, our data show that their respective demographic histories following colonisation were also distinct. On the one hand, Crete harbours a small population and maintains very low levels of gene flow with neighbouring populations. This has resulted in low genetic diversity, strong genetic drift, increased relatedness in the population and remote inbreeding. Cyprus, on the other hand, appears to maintain enough gene flow with the mainland to avoid such an outcome. Our work provides a comparative population genomic analysis of the effects of neutral processes on a classical island-mainland model system. It provides empirical evidence for the role of stochastic processes in determining the fate of diverging isolated populations.

## Introduction

Given their discrete borders, geographical isolation and abundance, islands are ideal systems to study patterns of genetic diversity in natural populations (Losos & Ricklefs, 2009). Due to the combination of biotic, abiotic, and stochastic forces, no two insular populations share the same demographic history (MacArthur & Wilson, 1967). Their fate is shaped by the timing of colonisation, fluctuations in population size and connectivity to neighbouring populations. These are directly impacted by the characteristics of the island, like carrying capacity and distance to the mainland, as well as the circumstances of colonisation such as bottlenecks and founder effects. The combined actions of reduced gene flow, *in situ* genetic drift, selection and potentially mutation influence the degree to which insular populations diverge (Barton, 1996; Grant, 1998; Mayr, 1954). Small populations are particularly sensitive to the effect of genetic drift, accelerating divergence from the surrounding populations. While high levels of gene flow can counter this effect, the lack of it can facilitate local adaptation by maintaining locally advantageous alleles (Tigano & Friesen, 2016) but can also lead to inbreeding with detrimental consequences (Frankham, 1998).

In small isolated populations, without other sources for genetic diversity besides mutation and recombination, the relatedness among insular individuals increases over time under the effect of drift. As a result, levels of remote inbreeding may rise even with the avoidance of mating between close relatives. Although this is a common occurrence in island populations, mating between related individuals can lead to inbreeding depression (Keller & Waller, 2002) and, in extreme circumstances, local extinction (Frankham, 1997, 1998). As such, the study of the genetic makeup of insular populations can provide key information from a conservation perspective. Despite being widely used to estimate inbreeding and infer demographic histories, traditional genetic markers lack resolution to reconstruct particularly convoluted systems such as, for example, multiple islands or among modestly differentiated populations. Technological advances now provide more affordable high-representation genomic data such as the sequencing of whole genomes. Combined with increasingly sophisticated methods, it allows for more accurate inferences, even for non-model species (Ellegren, 2014).

The eastern Mediterranean offers an excellent setting to study insular demographic history. A biodiversity hotspot (Médail & Quézel, 1999), the area is riddled with islands, the largest of which are Crete (CT) and Cyprus (CY). While fluctuating sea levels intermittently connected smaller islands to the mainland in the Quaternary, CT and CY have been isolated since the end of the Messinian salinity crisis (approx. 5 Mya.; Bache et al., 2012). They share many common features such as distance to mainland (95 and 75 km, respectively), surface area (8500 and 9200 km^2^) and a Mediterranean-subtropical climate with mild winters and warm summers. Their strategic position makes them pivotal stop-overs in the seasonal migration of many bird species, and movements of bird populations are widely studied (e.g. Emin et al., 2018; Panter et al., 2020). However, thus far they have been the subject of only few genetic studies, most on each island individually rather than comparatively, and typically focusing on human commensal small mammal species (Bonhomme et al., 2011; Cucchi, Vigne, Auffray, Croft, & Peltenburg, 2002; Dubey et al., 2007).

The Afro-European barn owl (*Tyto alba*) is a non-migratory bird of prey present across the African and European continents, as well as most of the surrounding islands and archipelagos (Uva, Päckert, Cibois, Fumagalli, & Roulin, 2018). In spite of being quite widespread and maintaining high gene flow overland (Antoniazza, Burri, Fumagalli, Goudet, & Roulin, 2010; Machado, Clément, Uva, Goudet, & Roulin, 2018), populations separated by water barriers appear to accumulate differentiation more quickly, with numerous insular subspecies (Burri et al., 2016; Machado et al., 2021; Uva et al., 2018). In the eastern Mediterranean, the continental European barn owl lineage meets the eastern subspecies *T. a. erlangeri* (W. L. Sclater, 1921) from the Levant (Burri et al., 2016; Cumer et al., 2021). Although Crete and Cyprus populations supposedly belong to *T. a. erlangeri* (Clements et al., 2019), the low resolution genetic data previously available was insufficient to clarify the history of each island and how they relate to the mainland. Barn owls from Crete appeared to be quite distinct from all surrounding mainland, including the Levant (Burri et al., 2016), and the demographic history of the Cyprus owl population has never been studied.

Here, we investigate the genetic structure and past demographic history of insular and mainland barn owl populations in the eastern Mediterranean. We focus in particular on Crete and Cyprus, the two largest islands in the region, that have very similar intrinsic characteristics and are thought to harbour barn owls from the eastern subspecies found in the Levant (*T. a. erlangeri*). As such, the populations should have originated, independently or not, from the Levant. However, being closer to other Greek islands and the Greek mainland, Crete could have actually been colonised from there, which would be incompatible with it belonging to the same subspecies as Cyprus. Taking advantage of the whole genome sequences of 65 individuals and the recent publication of a high-quality reference genome (Machado et al., 2021), we address this by modelling the colonisation of both islands from the mainland. Lastly, we compare how their different demographic histories impacted their current genetic diversity and inbreeding levels.

## Materials and Methods

### Sampling, Sequencing and Genotyping

A total of 67 barn owl individuals from seven populations were used in this study (Table 1; Supporting Table 1): 10 in Italy (IT), 5 in islands of the Ionian Sea (IO), 10 in Greece (GR), 11 in islands of the Aegean Sea (AE), 11 in Crete (CT), 10 in Cyprus (CY) and 10 in Israel (IS). Of these, 47 were sequenced in Cumer et al. (2021; GenBank BioProject PRJNA727977; Sup. Table 1). One additional individual of the Eastern barn-owl species (*T. javanica* from Singapore; Uva et al., 2018) was used as an outgroup for specific analyses. The outgroup was sequenced in Machado et al., (2021; GenBank BioProject PRJNA700797). The remaining 20 samples followed the same protocol described in (Cumer et al., 2021; Machado et al., 2021). In brief, we extracted genomic DNA using the DNeasy Blood & Tissue kit (Qiagen, Hilden, Germany) and prepared individually tagged 100bp “TruSeq DNA PCR-free” libraries (Illumina) following the manufacturer’s instructions. Then, whole-genome resequencing was performed on multiplexed libraries with Illumina HiSeq 2500 high-throughput paired-end sequencing technologies at the Lausanne Genomic Technologies Facility (GTF, University of Lausanne, Switzerland) with an expected sequence coverage of at least 15X.

**Table 1.** Population genetic diversity, inbreeding and divergence estimates for barn owls of the eastern Mediterranean. The standard deviations of the values are provided between brackets for each parameter except for population specific *F*_ST_ where values are the standard error of the mean.

| Pop | Abbr. | N | # PA | # PA lin. | # mtDNA | $\pi$ mtDNA | H<sub>O</sub> | Pop $F_{ST}$ | $F_{IS}$ | $F_{IT}$ | $F_{ROH}$ | $\beta$ |
| --- | --- | --- | --- | --- | --- | --- | --- | --- | --- | --- | --- | --- |
| Italy | IT | 9 | 118'152<br>(202) | 188'285 | 68<br>(23) | 0.0021<br>(0.0011) | 0.164<br>(0.002) | 0.058<br>(0.005) | -0.024<br>(0.012) | 0.014<br>(0.009) | 0.028<br>(0.01) | 0.112<br>(0.007) |
| Ionian Islands | IO | 5 | 46'340 |  | 87 | 0.0021<br>(0.0013) | 0.16<br>(0.004) | 0.091<br>(0.004) | -0.039<br>(0.027) | 0.019<br>(0.02) | 0.067<br>(0.02) | 0.143<br>(0.041) |
| Greece | GR | 10 | 73'108<br>(220) | 239'089<br>(294) | 86<br>(19) | 0.0021<br>(0.0012) | 0.165<br>(0.005) | 0.047<br>(0.003) | -0.02<br>(0.029) | 0<br>(0.022) | 0.038<br>(0.02) | 0.101<br>(0.008) |
| Aegean Islands | AE | 11 | 79'357<br>(198) |  | 81<br>(18) | 0.0023<br>(0.0012) | 0.164<br>(0.01) | 0.038<br>(0.002) | 0<br>(0.059) | 0.013<br>(0.049) | 0.043<br>(0.03) | 0.092<br>(0.014) |
| Crete | CT | 11 | 82'202<br>(177) | 124'440<br>(129) | 51<br>(24) | 0.0013<br>(0.0007) | 0.153<br>(0.006) | 0.115<br>(0.005) | -0.018<br>(0.037) | 0.05<br>(0.034) | 0.086<br>(0.04) | 0.165<br>(0.024) |
| Cyprus | CY | 10 | 121'550<br>(196) | 177'675<br>(113) | 72<br>(22) | 0.002<br>(0.001) | 0.165<br>(0.008) | 0.061<br>(0.005) | -0.032<br>(0.05) | 0.029<br>(0.039) | 0.04<br>(0.03) | 0.114<br>(0.024) |
| Israel | IS | 9 | 271'400<br>(235) | 413'375 | 43<br>(20) | 0.0013<br>(0.0007) | 0.175<br>(0.003) | 0.007<br>(0.003) | -0.035<br>(0.016) | 0.027<br>(0.008) | 0.019<br>(0.01) | 0.063<br>(0.012) |

N: number of individuals in the population; #PA: private alleles per population, bootstrapped to the smallest N of 5 individuals; #PA lin.: private alleles per lineage of K=5 identified with sNMF, bootstrapped to the smallest N of 9 individuals; #mtDNA: mitochondrial polymorphic sites per population, bootstrapped to the smallest N of 5 individuals; $\pi$ mtDNA: mitochondrial nucleotide diversity; H<sub>O</sub>: observed heterozygosity; $F_{ST}$ : population specific $F_{ST}$ as in (Weir and Goudet 2017) bootstrapped over 100 blocks of contiguous SNP; $F_{IS}$ : population level inbreeding coefficient; $F_{IT}$ : mean individual inbreeding coefficient relative to the meta-population; $F_{ROH}$ : mean inbreeding coefficient estimated from ROH; $\beta$ : mean pairwise relatedness within population.

The bioinformatics pipeline used to obtain analysis-ready SNPs from the raw sequenced of the 65 individuals plus the outgroup was the same as in (Machado et al., 2021) adapted from the Genome Analysis Toolkit (GATK) Best Practices (Van der Auwera et al. 2013) to a non-model organism following the developers’ recommendations. Briefly, we trimmed the reads to 70bp length with Trimommatic v.0.36 (Bolger et al. 2014) and aligned them with BWA-MEM v.0.7.15 (Li and Durbin 2009) to the barn owl reference genome (GenBank accession JAEUGV000000000; Machado et al., 2021). Then, we performed base quality score recalibration (BQSR) following the iterative approach recommended for non-model species that lack a set of “true variants” in GATK v.4.1.3 using high-confidence calls obtained from two independent callers: GATK’s HaplotypeCaller and GenotypeGVCF v.4.1.3 and ANGSD v.0.921 (Korneliussen et al. 2014). Following BQSR, we called variants with GATK’s HaplotypeCaller and GenotypeGVCFs v.4.1.3 from the recalibrated bam files.

For variant filtering we followed GATK hard filtering suggestions for non-model organisms, with values adapted to our dataset and expected coverage using GATK v4.1.3.0 and VCFtools v0.1.15 (Danecek et al., 2011). A detailed documentation of the filters applied can be found in Sup. Table 2. We also removed scaffolds that belong to the Z chromosome due to it being hemizygous in females (Sup. Table 1). In preliminary analyses we corrected the origin of a sample, an injured owl found at sea and reported to a port in mainland Greece but that was genetically of Cretan origin and considered as such hereafter. We also removed one Italian (IT10) and one Israeli (IS10) individuals as relatedness analyses revealed they were each part of a sibling pair. The final dataset contained 5’493’583 biallelic SNPs with a mean coverage of 16.4X (4.38 SD) across 65 individuals (Sup. Table 1).

### Mitochondrial DNA

#### Sequencing and assembly of mitochondrial genome

We produced a complete mitochondrial reference genome for the barn owl, from the same individual used for the reference nuclear genome recently published (Machado et al., 2021). The mitochondrial genome was thus produced from the high molecular weight (HMW) DNA extraction described in detail in (Machado et al., 2021). Briefly, HMW DNA was extracted from a fresh blood sample using the agarose plug method as described in (M. Zhang et al., 2012). Then, 15-20 kb DNA fragments were obtained with Megaruptor (Diagenode, Denville, NJ, USA) and checked on a Fragment Analyzer (Advanced Analytical Technologies, Ames, IA, USA). 5 µg of the sheared DNA was used to prepare a SMRTbell library with the PacBio SMRTbell Express Template Prep Kit 2.0 (Pacific Biosciences, Menlo Park, CA, USA) according to the manufacturer’s recommendations. The resulting library was size-selected on a BluePippin system (Sage Science, Inc. Beverly, MA, USA) for molecules larger than 13 kb. It was then sequenced on 1 SMRT cell 8M with v2.0/v2.0 chemistry on a PacBio Sequel II instrument (Pacific Biosciences, Menlo Park, CA, USA) at 30 hours movie length to produce HIFI reads.

After sequencing, we searched the circular consensus sequences (ccs) HIFI reads for sequences matching the 18128 bp mitochondrial genome of the previous assembly (NCBI Reference Sequence: NW_022670451.1; Ducrest et al., 2020) using minimap2 (Li, 2018) with the option -x asm5. We obtained twelve reads, which were reverse complemented as needed in order to be in the same orientation as our seed mitochondrial genome. No read was long enough to obtain a closed circular mitochondrial genome. Thus, we selected a css read of particularly high quality as an anchor and used two other overlapping reads to complete the circular sequence. From these three high quality reads, we manually assembled a full-length mitochondrial genome of 22461 bp. Mitochondrial css are provided in supplementary material and the reference sequence has been deposited at GenBank under the accession MZ318036.

We annotated the mitochondrial genome using MitoAnnotator v3.52 (Iwasaki et al., 2013) and removed the hyper-variable D-loop for the subsequent analyses, yielding a 15’571bp sequence.

#### Mitochondrial population structure and genetic diversity

To obtain the mitochondrial sequences of each individual, we mapped their trimmed whole-genome resequencing reads onto the newly assembled barn owl mitochondrial genome using the BWA-MEM v.0.7.15 algorithm (Li & Durbin, 2009). We then called variants using the bcftools v1.8 (Danecek et al., 2011) mpileup (with mapping quality > 60, depth < 5000) and call (consensus calling, -c) for haploid data (ploidy=1). We then created a consensus fasta sequence with bcftools consensus, applying variants called above on the reference genome. We aligned individual fasta sequences using ClustalOmega v1.2.4 (Sievers et al., 2011) and manually checked the alignment for errors in MEGA X v10.1.7 (Kumar, Stecher, Li, Knyaz, & Tamura, 2018). We generated a mitochondrial haplotype network using the R package pegas v0.14 (Paradis, 2010) and grouped similar haplotypes into haplogroups (Sup. Fig. 1). Finally, we quantified population diversity (nucleotide diversity, π) and divergence (*Φ*_ST_) with Arlequin v3.5.2.2 (Excoffier & Lischer, 2010).

### Population structure, diversity and inbreeding

To elucidate population structure in our dataset, we performed a principal component analysis (PCA) using the R-package SNPRelate v3.11 (Zheng et al., 2012) and inferred individual admixture proportions with the software sNMF v1.2 (Frichot, Mathieu, Trouillon, Bouchard, & François, 2014). sNMF was run for values of K ranging from 2 to 9, with 10 replicates for each K. Runs were checked visually for convergence within each K. For both analyses, we used a dataset of 603’496 biallelic

SNPs obtained by pruning our SNP dataset for linkage disequilibrium (LD) using PLINK v1.9 (--indep-pairwise 50 10 0.1; (Chang et al., 2015) as recommended by the authors. To investigate whether an island population was the product of admixture between two sampled populations, we used the f3 statistic (Patterson et al., 2012) and TreeMix (Pickrell & Pritchard, 2012) both calculated with the TreeMix v1.13 software. TreeMix was run in 20 replicates, using a bootstrap per 500 SNP interval, with 0 to 3 migration events, using the same LD-pruned dataset as above, to which any sites with missing data were removed yielding a total of 598’599 SNPs.

We used SNPRelate to calculate an allele sharing matrix between individuals (β; (Weir & Goudet, 2017) individual inbreeding coefficients relative to the total and then averaged per population (*F*_IT_). We used the R package hierfstat v.0.5-9 (Goudet, 2005) to estimate population pairwise and population-specific *F*_ST_ as in (Weir & Goudet, 2017). Confidence intervals were obtained by bootstrapping 100 times 100 blocks of contiguous SNPs. We also used hierfstat to quantify individual inbreeding coefficients relative to their population of origin and then averaged per population (*F*_IS_). For population genetic diversity, we calculated the observed individual observed heterozygosity and estimated the number of private alleles (i.e. alleles present in only one population) using custom made R scripts. To account for sample size differences in the estimation of private alleles, we subsampled 5 individuals (without replacement) from each population 100 times and calculated the mean number of private alleles in a population. When calculating the lineage-specific private alleles for K=5 from sNMF, we merged the populations of Greece, Ionian and Aegean islands and followed the same approach, this time sampling 9 individuals instead of 5 (corresponding to the new lowest sample size).

The Estimated Effective Migration Surface (EEMS) v.0.9 software (Petkova, Novembre, & Stephens, 2016) was used to visualize relative gene flow over the sampled region. First, we used the tool bed2diff to compute the matrix of genetic dissimilarities for the LD-pruned dataset mentioned above and utilized the Google Maps API v.3 tool (http://www.birdtheme.org/useful/v3tool.html) to draw a polygon outlining the study area. Then, EEMS was run with 700 demes in 3 independent chains of 2 million MCMC iterations with a 1 million iterations burn-in. We tested convergence of the results through a plot of observed-fitted values and the trace plot of the MCMC chain as suggested by the authors and plotted the results using the accompanying R package (rEEMSplots v.0.0.1).

We inferred runs of homozygosity (ROH) in the dataset by using the plink command --homozyg with default parameters (minimum 1 Mb length and 50 SNP). Only autosomal scaffolds of length more than 1 Mb were considered in ROH inference (47/70 scaffolds) covering 92% of the total assembly length. Given that bird chromosomes are typically shorter than those of humans (G. Zhang et al., 2014), for whom such methods were developed, we also called ROH with a minimum of 100Kb length. As the qualitative results were unchanged (data not shown), we kept the standard 1Mb threshold in a conservative approach to identify only identity by descent (IBD) segments and to facilitate potential comparisons with other studies. To estimate the index *F*_ROH_ we divided the sum of lengths of ROH in an individual with the length of the scaffolds (McQuillan et al., 2008) used after subtracting the number of ‘N’s (gaps) in the assembly. To visualize the distribution of ROH lengths per population, we divided ROH into five length classes: i) from 1Mb to under 2Mb, ii) from 2Mb to under 4Mb, iii) from 4Mb to under 6Mb, iv) from 6Mb to under 8Mb and finally, v) 8Mb or longer. We then calculated the number of base pairs falling within each ROH length class for every individual and averaged the values for each population.

To compare the levels of inbreeding, we tested whether *F*_IT_, *F*_ROH_ and β differ significantly between populations using a non-parametric Kruskal-Wallis rank sum test since the normality assumption did not hold. Further, we performed a pairwise Wilcoxon rank sum exact test with a Bonferroni correction for multiple testing to assess significance in the differences between pairs of populations. Given the small sample sizes (Table 1), we excluded obvious hybrid individuals (AE01, CT06) to avoid biasing the average of their respective populations.

### Demographic history

#### Demographic scenarios and parameters

To infer the origin and connectivity of the major insular barn owl populations (CT and CY), we used the software fastsimcoal2 (Excoffier, Dupanloup, Huerta-Sánchez, Sousa, & Foll, 2013). It uses coalescence simulations to estimate the composite likelihood of simulated demographic models under the observed site frequency spectrum (SFS). To model both island systems together, we would need to simulate the coalescence of the European and Levant lineages (sNMF K=2, Sup. Fig. 2>) for which we have no time calibrating event and could be hundreds of thousands of generations in the past. Such inference would likely be unreliable as well as extremely consuming computationally. Thus, we inferred the demographic history of each island system separately, including their closest populations. For each island system, ‘Crete’ and ‘Cyprus’, we tested three demographic scenarios (Figure 2b).

To infer the history of ‘Crete’, we did not include IS in the simulated scenarios as population structure analyses show that CT’s origin is not in the Levant, but rather from the European lineage (Figure 1). As such, we only considered the populations of AE and GR. The first two scenarios assume that both the Aegean islands and the island of Crete were colonized independently from the Greek mainland population. In the first one, the colonization of Crete takes place after the colonization of the Aegean islands, while in the second scenario Crete is colonized first. The third demographic scenario assumes the islands are colonized in a stepping-stone fashion, with owls from mainland Greece reaching the Aegean islands first and from there colonizing Crete (Figure 2b). Due to the low sea levels at Last Glacial Maximum (LGM), the Aegean islands were part of a larger emerged land mass that allowed nearly continuous overland connectivity to the mainland (Simaiakis et al., 2017). As such, for every demographic scenario in ‘Crete’ we assumed that the colonization of the Aegean islands from Greece occurred at the LGM (rounded to 18’000 years BP, 6’000 generations with a 3-year generation time). While the exact date is an approximation, allowing for migration between all populations after they split should reduce potential biases.

**Figure 1.**
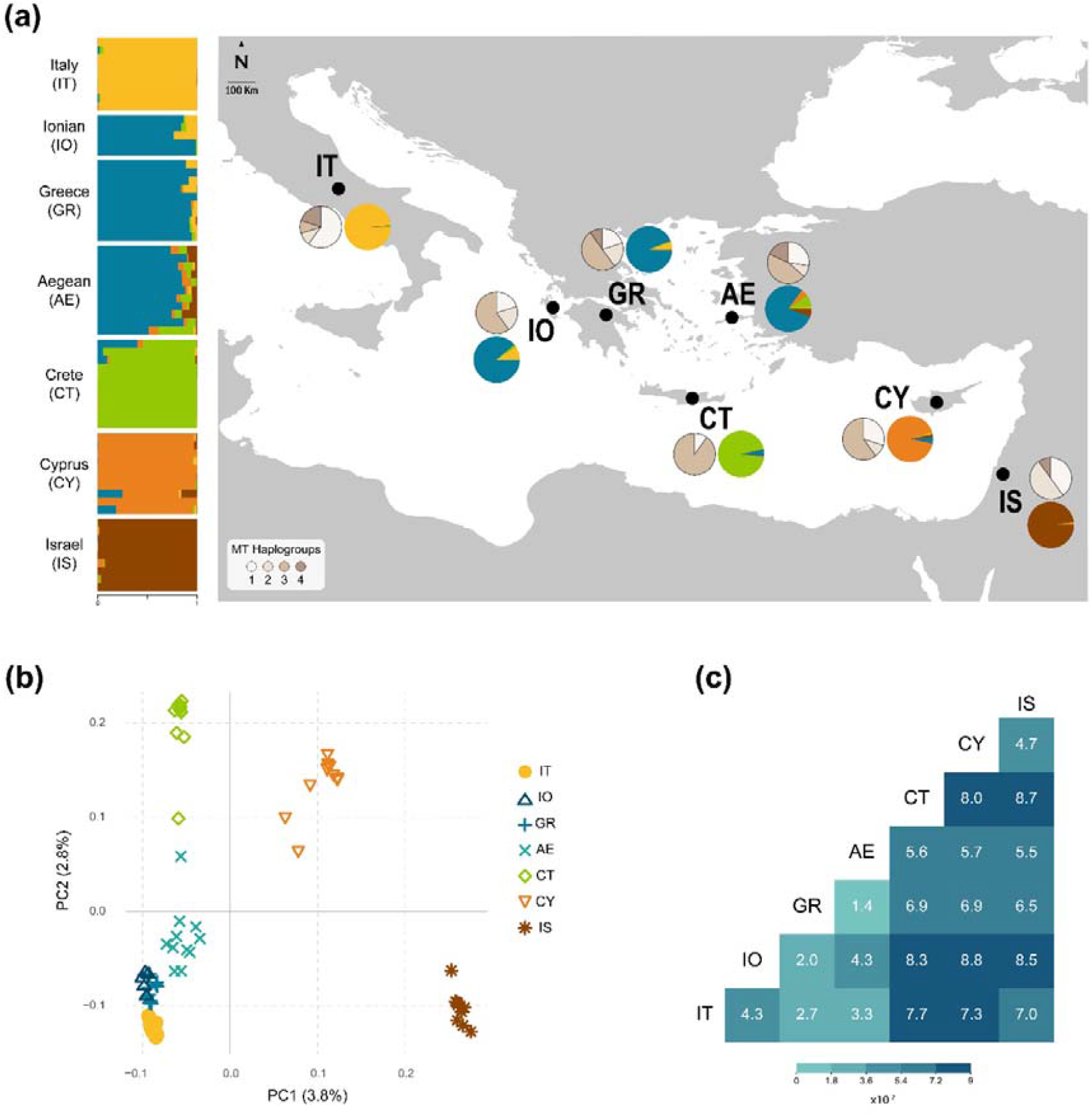
Population structure of barn owls in the Eastern Mediterranean. **(a)** Nuclear and mitochondrial population structure. Horizontal bars indicate individual admixture proportions for K=5 as determined by sNMF. Black dots on map indicate the approximate centroid of each population; coloured pie charts represent the mean admixture proportions per population; pie charts in shades of beige represent mitochondrial haplogroup proportions per population. **(b)** PCA based on the pruned nuclear SNP set. Values in parenthesis indicate the percentage of variance explained by each axis. **(c)** Pairwise FST between sampled barn owl populations. Heat map illustrates the given values according to the legend.

**Figure 2.**
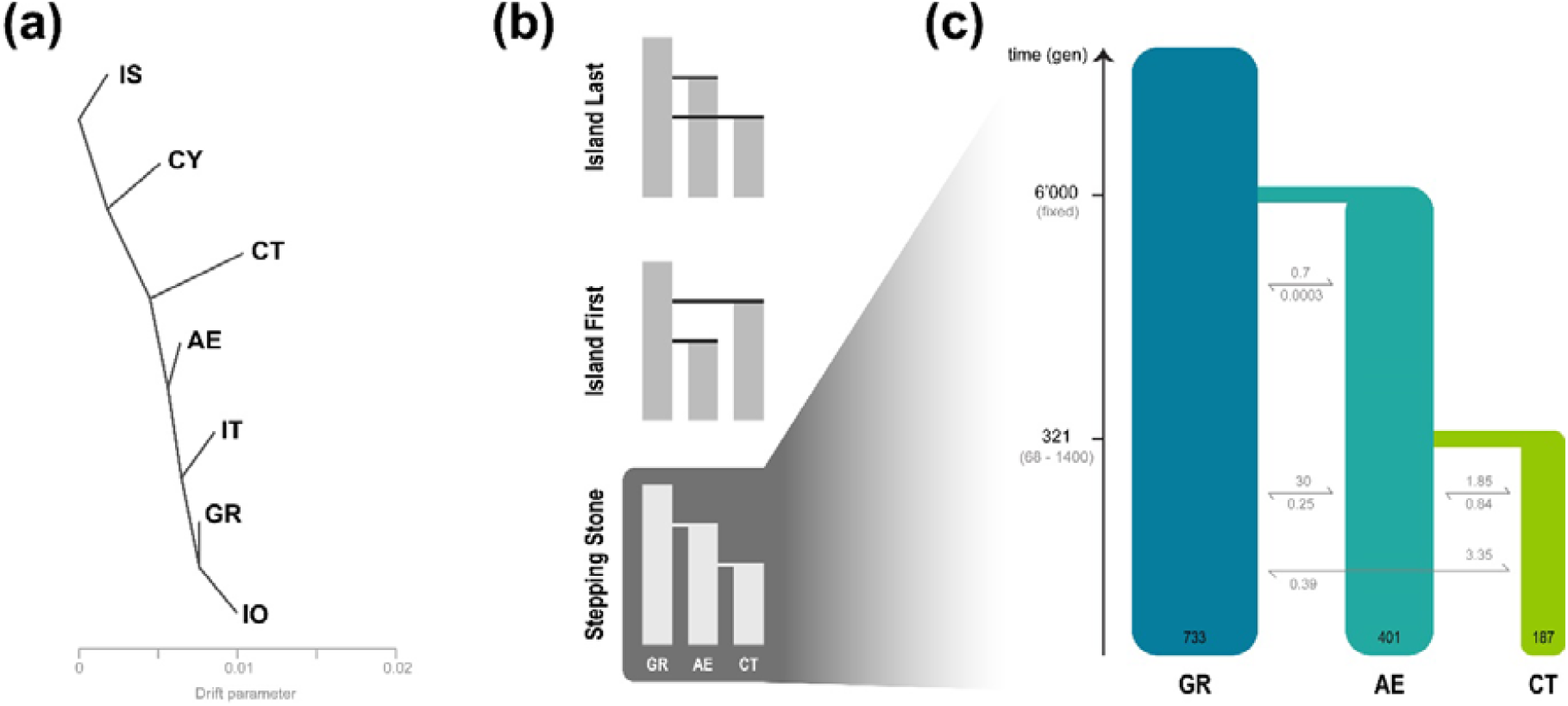
Demographic history of barn owl insular populations in the Eastern Mediterranean. **(a)** Treemix analysis with zero migration events. Population abbreviations follow Figure 1. **(b)** Hypothesized demographic topologies for the colonisation of Crete. The same topologies were tested for Cyprus, with IS instead of GR, “Ghost” instead of AE and CY instead of CT. **(c)** Best supported demographic model for the colonisation of Crete as determined by fastsimcoal2. Time is indicated in generations, confidence intervals at 95% are given between brackets. Population sizes (diploid) are shown at the bottom of each population bar; arrows indicate forward-in-time number of migrants per generation.

For ‘Cyprus’, in addition to IS as a representative of the Levant origin, a ghost population was incorporated in an attempt to represent the unsampled Turkish coast north of Cyprus, where the distance from the island to the mainland is the shortest. Including this ghost population in the model served two purposes. First, to account for unsampled sources of migrants into CY. Second, to avoid inflating artificially the effective population size of the CY population to justify the non-negligible admixture signal from AE (Figure 1a) that the simulator might interpret as *in situ* mutations. In the first two scenarios, both the Ghost and Cyprus populations originate from Israel, with the difference being the order in which they are colonized (same topology as Figure 2b). For the third scenario, owls from Israel would give origin to the Ghost population first and from there reach Cyprus.

#### Data preparation

Population sizes were reduced to the number of the smallest population in each model, resulting in 10 individuals per population for ‘Crete’ and 9 for ‘Cyprus’ (Sup. Table 1). To calculate the observed SFS for both systems, we filtered the data to a homogenous set of neutral markers. Specifically, we only kept sites with no missing data and with a depth of coverage less than 2/3 standard deviation from the mean. We also excluded CpG mutations (Pouyet, Aeschbacher, Thiéry, & Excoffier, 2018) and SNPs in genic regions. We inferred the ancestral state of the SNPs using the barn owl from Singapore, an outgroup to all our populations (Uva et al., 2018). Where the outgroup was homozygous for an allele, we marked that allele as the ancestral under rules of parsimony, while any other sites were removed. Population pairwise SFS were produced from the filtered datasets, giving 479’244 and 477’987 SNPs for ‘Crete’ and ‘Cyprus’, respectively.

#### Demographic inference with fastsimcoal2

For each system and each scenario, we specified a range of parameters from which the software drew an initial number as input in the optimization cycle (Sup. Table 4, 6). We modelled population splits with an instantaneous bottleneck in which the founding population size is a fraction of the present size.

For each scenario and each island, we performed 100 software runs. For each run we set the number of coalescent simulations to 500’000 and estimated the parameters through 50 expectation-maximization (EM) cycles. As we do not currently have a good estimation of the barn owl mutation rate, the end of the glaciation (rounded to 6000 generations ago) was fixed and all other parameters were scaled relative to it using the -0 option (based solely on polymorphic sites).

The best-fitting scenario was determined using Akaike’s information criterion (AIC; Akaike 1974). For the best scenario of each system, we performed non-parametric bootstrapping to estimate the 95% confidence intervals of the inferred parameters. Specifically, we divided the SNP dataset in 100 blocks with an equal number of SNPs, from which we created 100 bootstrapped-SFS and performed 50 independent runs of the software for each, with 250’000 simulations. Due to computational constraints we reduced the number of EM cycles to 10, an approach used previously and characterized as conservative (Malaspinas et al., 2016). The highest likelihood run for each bootstrapped replicate was used to calculate the 95% CI of the inferred parameters.

### Ancient population size inference

For inference of past effective population sizes, we used the Pairwise Sequential Markovian Coalescent (PSMC; Li and Durbin 2011). Specifically, we intended to estimate sizes in the distant past as this method is inaccurate for recent events. We ran the software on every individual of every population and calculated the median size for a population for each time interval. PSMC was executed with the same parameters as in (Nadachowska-Brzyska, Li, Smeds, Zhang, & Ellegren, 2015) (-N30 -t5 -r5 -p 4+30*2+4+6+10). For plotting we used a mutation rate of 8.28*10^−9^ mutations per site per generation as estimated for avian species by (Smeds, Qvarnström, & Ellegren, 2016) and a generation time of 3.6 years (Altwegg, Roulin, Kestenholz, & Jenni, 2006).

## Results

### Population structure and divergence in the eastern Mediterranean

Mitochondrial DNA exhibited an overall *Φ*_ST_ of 0.13 (AMOVA) across all sampled individuals and a range of nucleotide diversity (0.0013 – 0.0023; Table 1). The mitochondrial DNA analyses failed to show consistent population structure in the dataset. The first two haplogroups constructed from the haplotype network (Sup. Fig. 1) were present in all populations, while haplogroup 3 which was missing from Israel despite being predominant in nearby Cyprus, and haplogroup 4 which was found only on the mainland populations (Figure 1a). Cretan owls had the lowest haplogroup diversity with mostly haplogroup 3 present and the lowest nucleotide diversity (0.0013).

Principal component analysis based on whole nuclear genome SNP separated the populations approximately from West to East along the first axis with individuals for each population clustering together (Figure 1b), similar to K=2 in sNMF (Sup. Fig. 2). The second axis separated the two islands (CT & CY) from the rest of the populations, with admixed individuals dispersing between sources of admixture. Admixture analyses with sNMF were consistent between runs up to K=5 (Sup. Fig. 2, Figure 1a). For K=3, Crete separates from the European lineage and for K=4 CY separates from the Levant lineage (Sup. Fig. 2). For K=5 (Figure 1a), Italy, Crete, Cyprus, and Israel formed separate clusters while owls from the Ionian islands, mainland Greece and the Aegean were grouped into a single population. Owls from the Aegean islands showed the highest proportion of admixture (mean=0.2, SD=0.1) with components from Crete, Cyprus, and Israel in addition to their majority Greek component (Figure 1a). Some individuals from Crete and Cyprus appeared admixed between their respective island’s and the Greek component (blue in Figure 1a).

Tests for population admixture with f3 yielded a single slightly but significantly negative value (f3=- 0.00065, SE=6e-05, Z=-10.375), which showed the Greek population to be the product of admixture between the Aegean and the Ionian populations. None of the insular populations appeared to be the product of admixture between any population sampled in this study. The topology created by TreeMix was rooted at IS, with CY splitting first. CT displayed the longest branch of genetic drift and split before AE and the rest of the European populations (Figure 2a). The first migration event was from AE to GR (Sup. Fig. 3), and it was the only one consistent across runs.

Pairwise nuclear *F*_ST_ values ranged from 0.014 to 0.088 (Figure 1c), with the highest found between Cyprus and Ionian (0.088), followed by between Crete and Israel (0.087). Crete exhibited overall the highest pairwise values with any population (all above 0.056). Matching population divergence, the quantitative depiction of gene flow through EEMS identified a strong barrier to migration around the island of Crete and regions of reduced migration around the southern Ionian islands and the island of Cyprus (Sup. Fig 5).

### Genetic diversity and inbreeding

Genetic diversity based on nuclear SNP was generally highest in Israel and lowest in Crete, with Cyprus bearing comparable levels to any mainland population (Table 1). This was consistent for nuclear heterozygosity, population specific *F*_ST_ and gene diversity as well number of polymorphic sites in mtDNA. Private alleles were lowest among the closely related populations of Greece, the Ionian and Aegean Islands with Israel boasting the highest number. When considering GR, IO and AE as a genetic cluster (Figure 1a), Crete actually had the lowest number of private alleles (Table 1).

*F*_IS_, the average inbreeding coefficients of individuals relative to their population, was slightly negative in all populations (Table 1), as expected with random mating in a species with separate sexes (Balloux, 2004). A Cretan individual had a local inbreeding coefficient below -0.1, likely due to it being a F1 hybrid between CT and AE (see individual bars in Figure 1a and PC2 in Figure 1b) and two samples in the Aegean islands had a local inbreeding coefficient larger than 0.1 (Table 1; Figure 3b). *F*_IT_ values, the average inbreeding of individuals relative to the total set, but averaged per population, were highest on the island of Crete followed by the Aegean and Cyprus. Israel had significantly lower *F*_IT_ than all other populations, whereas Crete’s was higher than all but the Aegean (Figure 3a; Table 1; *X*^2^= 36.043, *p* < 0.001). Thus, Cyprus had higher F_IT_ than Israel, smaller than Crete and similar to every other population. Individual relatedness (β) was highest between two Ionian individuals found to be half-sibs (Sup. Fig. 4). Otherwise, individuals from Crete were more related to each other than any other pair of individuals in the dataset (Table 1; *X*^2^= 195.77, *p* < 0.001). In its turn, individuals from Cyprus were only more related to each other on average than the populations of Israel and the Aegean.

**Figure 3.**
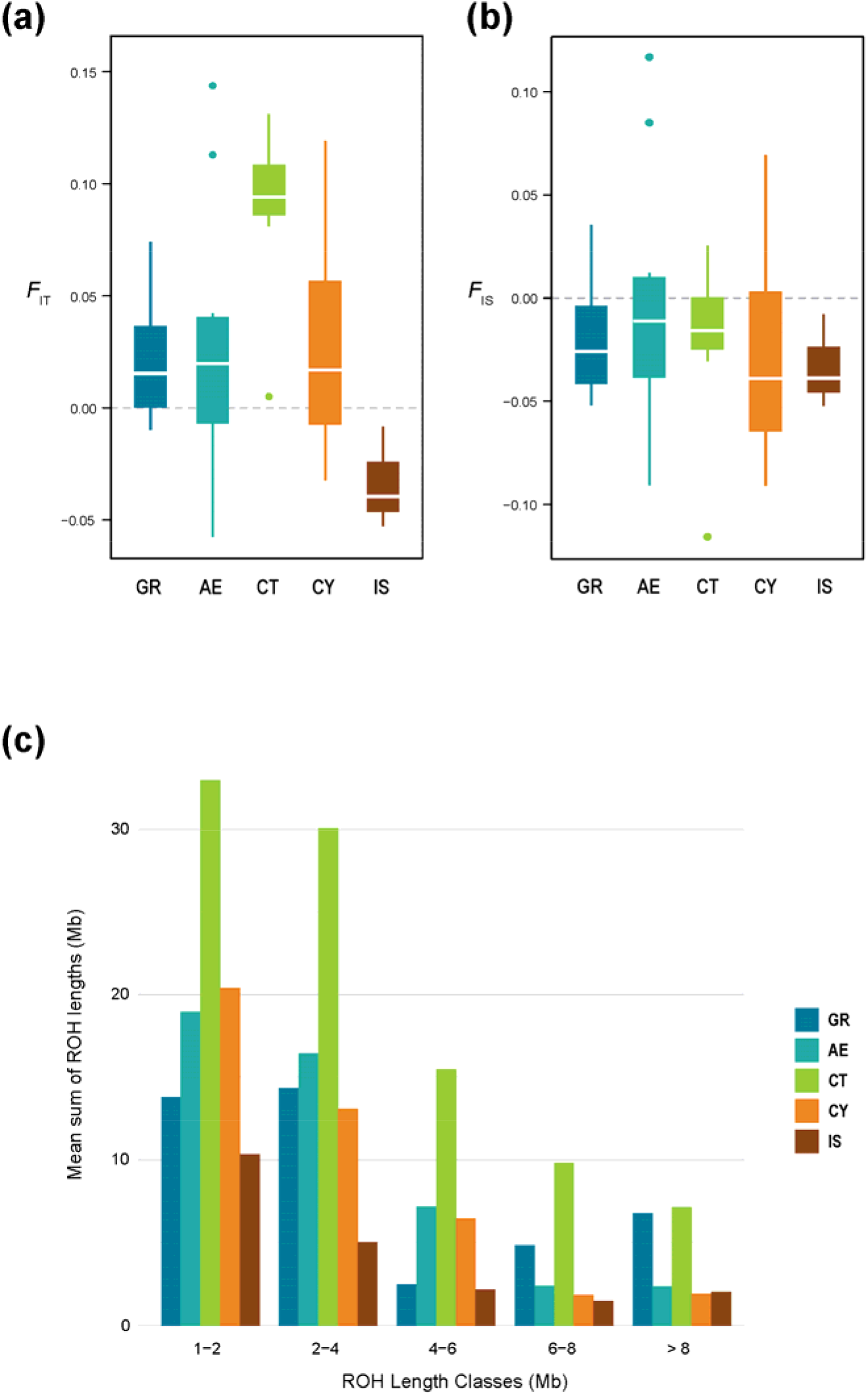
Comparison of inbreeding in insular barn owls to their closest mainland counterparts in the Eastern Mediterranean. (a) *F*_IT_ measure of inbreeding calculated from individual allele matching proportions relative to the average in the dataset (dashed line is *F*_IT_=0). (b) *F*_IS_ measure of inbreeding calculated from individual allele matching proportions relative to the average in the subpopulation (dashed line is FIS=0). (c) Per population average length of ROH segments (in Mb) in each ROH length class.

Mean population *F*_ROH_ (i.e. proportion of the genome in runs of homozygosity) were also highest in Crete, followed by the Ionian, Aegean and Cyprus (Table 1). Individuals from Crete showed the highest proportion of ROH of all sizes (Figure 3c; Sup. Fig. 6), while individuals from Israel had the lowest proportion in all categories. Individuals from Cyprus and the Aegean were also enriched in ROH segments compared to their mainland origin in most length classes, but much less so than Crete (Figure 3c). Indeed, while *F*_ROH_ was significantly higher in Crete than in Greece, it was not the case between the Aegean and Greece nor between Cyprus and Israel (*X*^2^= 11.862, *p* < 0.001).

### Demographic history

We simulated three different demographic scenarios for each island system, two where the island was colonized from the mainland either before or after the other population in the model (AE for “Crete” and Ghost for “Cyprus”) and one where the populations are colonized in a stepping-stone manner (Figure 2b). The best demographic scenario inferred with fastsimcoal2 for the island of Crete was the stepping-stone model (Figure 2c; Sup. Table 3). Here, the Cretan population originates very recently from the Aegean islands 321 generations BP (68-1400 95% CI), itself colonized from mainland Greece at the fixed time of 6000 generations BP (Sup. Table 5). Estimated migration rates were higher towards the island from both GR and AE (6.7 and 3.7, respectively) and lower in the other direction (0.7 and 1.7). Inferred effective population sizes were highest for the Greek (1465 haploids; 509-7880 95% CI) mainland and lowest for Crete (373 haploids; 107-944 95% CI). Past instantaneous bottlenecks at colonisation were pronounced both for the Aegean and Cretan populations (48 [13-2922 95% CI] and 74 [6-243 95% CI] haploids, respectively).

For the island of Cyprus, the best-fitting scenario consisted of colonization from Israel after the colonization of the ‘Ghost’ population coinciding with a hypothesized mainland population residing on the southern coast of Turkey (similar topology as Figure 2b; Sup. Table 3). Colonization time for CY was much more recent than the last glaciation (986 generations, less than 3’000 years BP; Sup. Table 7). However, the Ghost population was estimated to have an unrealistic large effective population size (65’310 diploids), and CY an extremely small one (61). The migration rates inferred indicate a complete replacement of CY each generation by the Ghost, suggesting this model is far from being an accurate representation of reality. As such we interpret its results with caution.

PSMC identified a pronounced bottleneck for all populations (around 20’000 years BP) but failed to show a clean split for the two Islands, particularly Crete, and any mainland population (Sup. Fig. 7).

## Discussion

Although insular populations have greatly contributed to the development of evolutionary theory (Grant, 1998; MacArthur & Wilson, 1967; Warren et al., 2015), the study potential of many of these remains untapped. The colonisation and settlement of an island by a given organism depend not only on the geographic context and specific island characteristics but also on stochastic events. As such, seemingly identical islands may yield populations with contrasting fates. Here, we investigate the demographic history and current patterns of inbreeding and genetic diversity of insular barn owls in the eastern Mediterranean Sea. In particular, we are interested in owls from Crete and Cyprus which, alongside the Levant region, were thought to form a subspecies *Tyto alba* erlangeri. These two similar islands in terms of size, climate and distance to mainland provide natural replicates for a comparative analysis of the colonisation and ensuing demographic processes. Using whole genome sequences, we show how each island and archipelago have unique histories and exhibit different degrees of isolation and the effect this has on the genomes of individuals. Specifically, Crete and Cyprus were colonized from distinct mainland locations, each from a different ancestral lineage, inconsistent with them belonging to the same subspecies. The population in Crete originated from the European lineage, more precisely from the Aegean islands, while the population in Cyprus came from the Levant in the east. Additionally, Crete underwent stronger genetic drift and inbreeding than Cyprus, resulting in a smaller and less diverse population.

### Insular populations in the eastern Mediterranean

In the broader context of the Western Palearctic, our study targets two islands in the region where the European and eastern lineages of barn owls meet (Cumer et al., 2021). This is clearly shown in the genomic PCA, where the mainland populations of Italy and Greece in southern Europe were opposed to that of Israel in the Levant, with insular populations placed along this west-to-east genetic gradient roughly according to their geographic position (Figure 1b, Sup Fig K=2). The main islands of Crete and Cyprus are the most genetically distinct populations (Figure 1a,c), consistent with previous results for Crete (Burri et al., 2016). Conversely, the Greek archipelagos – Ionian and Aegean – were genetically very similar to the Greek mainland population (Figure 1a,c) suggesting they remain highly connected genetically. Such patterns of genetic differentiation reflect the geographical isolation of CT and CY, in contrast to the Aegean and Ionian archipelagos that are closer to the mainland through a network of adjacent islands and islets.

Overall, our results confirm that water bodies are strong barriers to barn owl movement (Cumer et al., 2021; Machado et al., 2021). For example, distant populations in the mainland, such as GR and IT, were much more similar to each other than any insular population, regardless of how distant each of them are (Figure 1). Nonetheless, all insular populations showed small signals of admixture with their neighbouring populations (Figure 1a). This likely reflects the intricate geographic setting, as well as the overall low differentiation within this species (overall *F*_ST_ in our dataset 0.03, and 0.047 in the whole Western Palearctic (Cumer et al., 2021), that mtDNA data lacked the resolution to detect (Figure 1a, Table 1, Sup. Fig. 1; Burri et al., 2016). Insular populations had generally lower levels of population private diversity, while displaying similar levels of heterozygosity (Table 1), and higher within-population relatedness compared to the mainland (Sup. Fig. 4), reflecting their isolation. However, despite all populations appearing to mate randomly within localities (*F*_IS_ slightly negative as is expected from a dioecious species (Balloux, 2004); Figure 3b, Table 1), the inbreeding levels of insular barn owls relative to the whole set of populations were quite large (*F*_IT_ and *F*_ROH_; Figure 3a,b, Table 1).

### Crete and Cyprus

Despite the inherent physical similarities between the islands of Crete and Cyprus, their barn owl populations differ in many aspects. These natural replicates of island-mainland comparisons, with similar climatic conditions, supposedly harbour the subspecies *T. a. erlangeri* from the Levant (W. L. Sclater, 1921). However, while Cyprus’ genetically closest mainland population is indeed Israel, Crete is actually most similar genetically to Greece (Figure 1b). This demonstrates that Crete is not home to barn owls of the eastern subspecies, but rather from the European mainland lineage (*T. a. alba*). Not only they have separate geographic origins, but we also show that their *in situ* demographic histories are quite distinct.

Since its colonisation and founding bottleneck, Crete maintained a low population size with little gene flow with neighbouring populations (Figure 2c; Sup. Fig. 5), generating background relatedness among individuals (Sup. Fig. 4). The low gene flow it maintains with the surrounding populations (Sup. Fig. 5) may be due to the very strong winds that surround the island (Zecchetto & De Biasio, 2007) acting as a barrier by hindering flight. Thus, despite random mating within the island (low *F*_IS_), remote inbreeding increased (high *F*_IT_ and *F*_ROH_) due to high relatedness (high β), making CT the most inbred population by far in our dataset, as well as the least diverse (Table 1; Figure 3a,b). Accordingly, it carried the highest proportion of ROH compared to any other populations (Table 1). Notably, CT was enriched in ROH of all sizes (Figure 3c), suggesting a small effective size over a long time period until today (Ceballos, Joshi, Clark, Ramsay, & Wilson, 2018). This strong isolation coupled with small population size resulted in a very distinct genetic composition through the effect of genetic drift (Figure 2a) as well as high individual relatedness and inbreeding.

In contrast, Cyprus appears to have maintained enough gene flow with the mainland preventing it from accumulating remote inbreeding, while allowing for differentiation. Winds in this region are weaker than around Crete (Zecchetto & De Biasio, 2007), potentially facilitating the contact between Cyprus and Israel in the Levant, the most diverse population in our study. This could explain the surprisingly similar patterns of genetic diversity in CY to that of mainland populations (Table 1), which suggest a higher effective population size in spite of the inference from fastsimcoal2 (Sup. Table 7). Furthermore, CY had considerably less runs of homozygosity (ROH) than CT, carrying only a slight enrichment in short length classes, similar to the Aegean and Ionian islands (Figure 3c). Given the high inter-individual variability in relatedness and inbreeding coefficients (Table 1; Fig. 3a; Sup. Fig. 4), it appears that the gene flow with Israel and/or an unknown, unsampled population prevents the rise of population-wide inbreeding as observed in Crete. Interestingly, the most common mitochondrial haplogroup in CY was found in European populations but absent in IS (haplogroup 3, Figure 1a). Although it could simply have been unsampled in the Levant, it may also be evidence of some gene flow between the European and eastern lineages as seen in the two admixed individuals of CY (Figure 1a; see also paragraph after next)

Overall, the different levels of connectivity (i.e. levels of gene flow) of each island appear to be the main driver of their diverging histories. However, insular specificities may also contribute to this effect. The carrying capacity of CT and CY for barn owls could be different due to cryptic differences in nesting or roosting site availability, for example, in spite of their similar surface area. In addition, the mountainous landscape in CT could restrict dispersal movements as well as reduce the suitable surface for breeding. Finally, intrinsic characteristics of the colonisation of both islands may also have contributed to their diverging histories.

On the one hand, CY was colonised directly from the highly diverse and large mainland population of IS (Sup. Table 7). As such, both the settlers of the island and subsequent immigrants were likely unrelated and diverse, preventing the insular population from increasing steeply in relatedness. Our simulations suggest that colonisation occurred about 3000 years BP (1900 – 10000 years BP). However, this result should be interpreted cautiously as the modelling for this island system yielded unreasonable population size estimates (Sup. Table 7) likely due to our use of a ghost population to represent mainland Turkey. This is suspected to be a contact zone between the European and eastern barn owl lineages with sporadic gene flow (Cumer et al., 2021). Our observations support this hypothesis as islands on both sides of Turkey, namely CY and AE, carried some small genetic components from the other (Figure 1a). In this context, our modelled ghost population would likely be admixed or even outbred which would explain its exaggerated population size. Sampling in Turkey will be key to clarify this hypothesis and fully describe the dynamics between barn owl populations in the eastern Mediterranean.

On the other hand, demographic simulations showed that CT was colonised from the AE archipelago rather than directly from mainland Greece (Figure 2c; Sup. Table 5). This was supported by the second axis in the PCA which placed individuals in a gradient from GR to AE and then CT (Figure 1b). Remarkably, one AE owl from the south-eastern island of Rhodes had approximately 50% Cretan origin hinting at how the patchwork of islands and islets in the region could have been used as stepping stones during colonisation. Thus, CT was colonised from what is already a less diverse insular population, which in turn came from the GR mainland, itself less diverse than Israel (Table 1). This cumulative loss of diversity through recurrent bottlenecks and possible expansion could contribute to the quick increase in relatedness in the island given its small population size, despite its recent colonisation. Indeed, CT was inferred to have been colonised by barn owls around 1000 years BP (204 – 4200 years BP; Figure 2c; Sup. Table 5). Accordingly, PSMC failed to uncover any signal of older divergence (Sup. Figure 7). Nonetheless, considering the geological age of the island (5 million years BP) and that agricultural practices have been established there for millennia (Greig & Warren, 1974), this estimation appears extraordinarily recent. Absent any other source of evidence, one can only speculate as to why this population is so recent. It is possible that a massive migration led to population replacement at a time when sea levels were lower and the surrounding islands closer, masking any trace of an earlier settlement. Alternatively, earlier settlers could have been extinct due, for example, to a natural disaster such as the catastrophic Minoan volcanic eruption (3’500 years BP) (Pareschi, Favalli, & Boschi, 2006).

## Conclusion

Our work provides a comparative study on two natural replicates of island colonisation by the barn owl, a bird that despite being found in many islands avoids flying over open bodies of water. The use of whole genome sequences allowed us to demonstrate that Crete and Cyprus owls come from different genetic backgrounds, as each island originates from a distinct continental genetic lineage (Figure 1). Further, their histories diverge resulting in noticeably different populations. Cyprus was colonised directly from the most diverse mainland population, accumulated differentiation but also remained sufficiently connected with it to maintain high levels of genetic diversity and prevent inbreeding (Table 1, Figure 3). Crete was reached by island hopping in the Aegean from a less diverse mainland population. The small size and isolation of this island population facilitated the impact of genetic drift which, along with inbreeding, led to it diverging considerably from its founders despite the recent colonisation (Table 1, Figure 2). Although further analyses would be necessary to study the functional consequences of inbreeding in Crete, this study shines a light on a real-life illustration of stochasticity in the classical island-mainland model systems.

## Acknowledgements

We are grateful to the following institutions and individuals that provided samples or aided in sampling to our study: The European Barn Owl Network, Burke Museum of Natural History and Culture (Washington, USA), Guillaume Dumont, Sylvain Antoniazza and Reto Burri. We thank Céline Simon for her valuable assistance with molecular work, and Christian Iseli for the preparation and maintenance of our bioinformatics tools and databases. This study was funded by the Swiss National Science Foundation with grants 31003A-138180 & 31003A_179358 to JG and 31003A_173178 to AR.

## Author contributions

APM, AT, AR, JG designed this study; APM produced whole-genome resequencing libraries; AT, APM conducted the analyses with input from TC, EL; PB; VB, MC, NK, PL, FM provided samples; ALD, MD produced the mitochondrial reference genome and NG assembled it; APM, AT led the writing of the manuscript with input from all authors.

## Data availability

The raw Illumina reads for the whole-genome sequences produced here have been deposited at GenBank BioProject PRJNA727915. In addition, we used date from BioProjects PRJNA727977 and PRJNA700797.

