## Supplementary Material for "Genomic consequences of colonisation, migration and genetic drift in barn owl insular populations of the eastern Mediterranean"

**Supplementary Tables**

**Supplementary Table 1** – Description of samples used in this study. Individuals retained for the inference with fastsimcoal2 are indicated with ^†^. Individuals marked with ^‡^ were removed from the analyses as they were each part of a sibling pair.

| **#** | **Pop** | **ID** | **Country** | **Location** | **Year** | **Tissue** | **Sex** | **Ref** |
| --- | --- | --- | --- | --- | --- | --- | --- | --- |
| 1 | Italy | IT01 | Italy | Roma | 2011 | blood | Female | 1 |
| 2 | Italy | IT02 | Italy | Roma | 2015 | blood | Female | 1 |
| 3 | Italy | IT03 | Italy | Roma | 2016 | blood | Male | 1 |
| 4 | Italy | IT04 | Italy | Roma | 2016 | blood | Male | 1 |
| 5 | Italy | IT05 | Italy | Roma | 2009 | blood | Female | 1 |
| 6 | Italy | IT06 | Italy | Grosseto | 2014 | blood | Female | 1 |
| 7 | Italy | IT07 | Italy | Livorno | 2001 | blood | Female | 1 |
| 8 | Italy | IT08 | Italy | Firenze | 2011 | blood | Female | 1 |
| 9 | Italy | IT09 | Italy | Firenze | 2016 | blood | Male | 1 |
| 10 | Italy | IT10^‡^ | Italy | Firenze | 2017 | blood | Female | 2 |
| 11 | Ionian | IO01 | Greece | Kefalonia Island | 2014 | soft tissue | Male | 2 |
| 12 | Ionian | IO02 | Greece | Kerkyra island | 2013 | soft tissue | Female | 2 |
| 13 | Ionian | IO03 | Greece | Kerkyra island | 2014 | soft tissue | Female | 2 |
| 14 | Ionian | IO04 | Greece | Zakynthos island | 2015 | blood | Male | 2 |
| 15 | Ionian | IO05 | Greece | Zakynthos island | 2015 | blood | Female | 2 |
| 16 | Greece | GR01^†^ | Greece | Agrinio | 2014 | blood | Male | 1 |
| 17 | Greece | GR02^†^ | Greece | Athens | 2014 | blood | Male | 1 |
| 18 | Greece | GR03^†^ | Greece | Chalandri | 2012 | blood | Male | 1 |
| 19 | Greece | GR04^†^ | Greece | Corinth | 2015 | blood | Female | 1 |
| 20 | Greece | GR05^†^ | Greece | Stoupa | 2015 | soft tissue | Female | 2 |
| 21 | Greece | GR06^†^ | Greece | Lamia | 2014 | soft tissue | Female | 1 |
| 22 | Greece | GR07^†^ | Greece | Mesolonghi | 2014 | soft tissue | Male | 1 |
| 23 | Greece | GR08^†^ | Greece | Morfovouni | 2015 | blood | Female | 1 |
| 24 | Greece | GR09^†^ | Greece | Panetolio | 2014 | blood | Female | 1 |
| 25 | Greece | GR10^†^ | Greece | Spata | 2015 | blood | Male | 1 |
| 26 | Aegean | AE01 | Greece | Rhodes island | 2014 | soft tissue | Female | 1 |
| 27 | Aegean | AE02^†^ | Greece | Rhodes Island | 2014 | soft tissue | Female | 1 |
| 28 | Aegean | AE03^†^ | Greece | Chios island | 2015 | blood | Male | 1 |
| 29 | Aegean | AE04^†^ | Greece | Chios island | 2012 | blood | Female | 1 |
| 30 | Aegean | AE05^†^ | Greece | Leros island | 2012 | blood | Male | 1 |
| 31 | Aegean | AE06^†^ | Greece | Lesvos Island | 2012 | blood | Male | 1 |
| 32 | Aegean | AE07^†^ | Greece | Lesvos Island | 2013 | soft tissue | Female | 1 |
| 33 | Aegean | AE08^†^ | Greece | Rhodes island | 2013 | blood | Male | 1 |
| 34 | Aegean | AE09^†^ | Greece | Leros island | 2014 | blood | Male | 1 |
| 35 | Aegean | AE10^†^ | Greece | Rhodes Island | 2015 | blood | Female | 1 |
| 36 | Aegean | AE11^†^ | Greece | Andros island | 2015 | blood | Male | 2 |
| 37 | Crete | CT01^†^ | Greece | Crete | 2015 | soft tissue | Male | 2 |
| 38 | Crete | CT02^†^ | Greece | Crete | 2015 | soft tissue | Male | 2 |
| 39 | Crete | CT03^†^ | Greece | Crete | 2015 | blood | Male | 2 |
| 40 | Crete | CT04^†^ | Greece | Crete | 2015 | soft tissue | Male | 2 |
| 41 | Crete | CT05^†^ | Greece | Crete | 2015 | blood | Male | 2 |
| 42 | Crete | CT06 | Greece | Crete | 2015 | blood | Female | 2 |
| 43 | Crete | CT07^†^ | Greece | Crete | 2013 | soft tissue | Female | 2 |
| 44 | Crete | CT08^†^ | Greece | Crete | 2014 | blood | Male | 2 |
| 45 | Crete | CT09^†^ | Greece | Crete | 2014 | blood | Female | 2 |
| 46 | Crete | CT10^†^ | Greece | Crete | 2014 | blood | Female | 2 |
| 47 | Crete | GR11^†^ | Greece | Kalamata | 2012 | blood | Female | 2 |
| 48 | Cyprus | CY01^†^ | Cyprus | Cyprus | NA | soft tissue | Female | 1 |
| 49 | Cyprus | CY02^†^ | Cyprus | Cyprus | 2016 | soft tissue | Female | 1 |
| 50 | Cyprus | CY03^†^ | Cyprus | Cyprus | 2016 | soft tissue | Male | 1 |
| 51 | Cyprus | CY04^†^ | Cyprus | Cyprus | NA | soft tissue | Female | 1 |
| 52 | Cyprus | CY05^†^ | Cyprus | Cyprus | NA | soft tissue | Male | 1 |
| 53 | Cyprus | CY06^†^ | Cyprus | Cyprus | NA | soft tissue | Female | 1 |
| 54 | Cyprus | CY07^†^ | Cyprus | Cyprus | 2017 | soft tissue | Female | 1 |
| 55 | Cyprus | CY08^†^ | Cyprus | Cyprus | 2015 | soft tissue | Female | 1 |
| 56 | Cyprus | CY09^†^ | Cyprus | Cyprus | 2017 | soft tissue | Female | 1 |
| 57 | Cyprus | CY10 | Cyprus | Cyprus | 2018 | soft tissue | Female | 1 |
| 58 | Israel | IS01^†^ | Israel | Lachish | 2005 | blood | Female | 1 |
| 59 | Israel | IS02^†^ | Israel | Beit Shean | 2005 | blood | Male | 1 |
| 60 | Israel | IS03^†^ | Israel | Hula | 2005 | blood | Female | 1 |
| 61 | Israel | IS04^†^ | Israel | Beit Shean | 2005 | blood | Female | 1 |
| 62 | Israel | IS05^†^ | Israel | Beit Shean | 2005 | blood | Male | 1 |
| 63 | Israel | IS06^†^ | Israel | Beit Shean | 2005 | blood | Female | 1 |
| 64 | Israel | IS07^†^ | Israel | Beit Shean | 2005 | blood | Female | 1 |
| 65 | Israel | IS08^†^ | Israel | Hula | 2005 | blood | Female | 1 |
| 66 | Israel | IS09^†^ | Israel | Hula | 2005 | blood | Female | 1 |
| 67 | Israel | IS10^‡^ | Israel | Hula | 2005 | blood | Female | 2 |
| 68 | Outgroup | SGP | Singapore | Singapore | 2015 | soft tissue | Female | 3 |

[1] Cumer et al. 2021 – GenBank BioProject PRJNA727977

[2] This study – GenBank BioProject PRJNA727915

[3] Machado et al. 2021 – GenBank BioProject PRJNA700797

**Supplementary Table 2**- Filters applied to raw variant calls. In the last column x denotes the values of the sites kept in the dataset.

| Parameter | Abbreviation in VCF file | Retained values |
| --- | --- | --- |
| Quality by Depth | QD | 5 < x |
| Root Mean Square Mapping Quality | MQ | 40 < x < 70 |
| Mapping Quality Rank Sum Test | MQRankSum | -12.5 < x |
| Read position Rank Sum Test | ReadPosRankSum | -8.0 < x |
| Strands Odds Ratio | SOR | x < 3.0 |
| Fisher strand | FS | x < 60 |
| Excess Heterozygosity | ExcessHet | x < 20 |
| Inbreeding coefficient | InbreedingCoeff | x < 0.9 |
| Site Depth | DP | 500 < x < 1600 |
| Genotype Depth | DP | 10 < x < 40 |
| Genotype Quality (Phred score) | GQ | 20 < x |
| Percent of missing data per site |  | x < 5% |
| HWE exact test (p-value) |  | 0.05 < x |

**Supplementary Table 3** – Likelihood and AIC comparison between simulated scenarios for systems Crete and Cyprus. Three topologies were tested for each system (Figure 2). The best scenario for each system is highlighted in bold.

| **System** | **Scenario** | **Likelihood** | **Δ likelihood** | **AIC** | **Δ AIC** |
| --- | --- | --- | --- | --- | --- |
| **CRETE** | A | -2’506’864 | 5’219 | 5’013’755 | 603 |
|  | B | -2’508’348 | 6’703 | 5’016’724 | 3’572 |
|  | **C** | **-2’506’562** | **4’917** | **5’013’152** | **0** |
| **CYPRUS** | **A** | **-878’596** | **-1’025** | **1’757’220** | **0** |
|  | B | -878’826 | -1’277 | 1’757’679 | 460 |
|  | C | -878’794 | -1’246 | 1’757’617 | 397 |

Likelihood – Maximum-likelihood estimated for the simulated SFS per demographic model; Δ likelihood – difference between the likelihood of the simulated and observed SFS; Δ AIC – delta AIC

**Supplementary Table 4**– Parameter ranges and estimates of fastsimcoal2 for system “Crete”. Initial range is the range from which the software draws initial the parameter values. The distributions are uniform for every parameter except *Bottleneck Intensity* which is log-uniform. Highest likelihood estimates are provided for every parameter. In scenarios A, C the Aegean islands diverge first from mainland Greece and therefore time of divergence for Crete is scaled to be less than 6’000 generations, and ancient migration is only between GR and AE. For scenario B, Crete is more ancient than AE and the time of split is ≥ 1 and ancient migration rates are between GR and CT. GR: Greece AE: Aegean CT: Crete.

| Parameter | Initial Range | Point estimates | | |
| --- | --- | --- | --- | --- |
|  |  | **Crete A** | **Crete B** | **Crete C** |
| Haploid population sizes | |  |  |  |
| *GR* | 500 – 100’000 | 502 | 7156 | 1465 |
| *AE* | 50 – 100’000 | 283 | 638 | 802 |
| *CT* | 10 – 4’000 | 162 | 1468 | 373 |
| Bottleneck Intensity (fraction of current population size) | | | | |
| *AE* | 0.01 – 0.5 | 0.187 | 0.064 | 0.060 |
| *CT* | 0.01 – 0.5 | 0.206 | 0.070 | 0.198 |
| Time of divergence (proportion of 6000; generations) | | | | |
| *T^CRETE^ (scenario A, C)* | 0.001 – 0.999 | 0.039 | - | 0.053 |
| *T^CRETE^ (scenario B)* | 1 – 3 | *-* | 1.218 | *-* |
| Ancestral migration rates (backwards in time) | | | | |
| *GR → AE (scenario A, C)* | 0 – 0.05 | 1e-06 | - | 1e-06 |
| *AE → GR (scenario A, C)* | 0 – 0.05 | 0.0018 | - | 0.0009 |
| *GR → CT (scenario B)* | 0 – 0.05 | - | 0.0029 | - |
| *CT → GR (scenario B)* | 0 – 0.05 | - | 0.0009 | - |
| Current migration rates (backwards in time) | | | | |
| *GR → AE* | 0 – 0.05 | 0.0332 | 0.0002 | 0.0006 |
| *AE → GR* | 0 – 0.05 | 0.0678 | 0.0429 | 0.0417 |
| *GR → CT* | 0 – 0.05 | 0.0005 | 0.0003 | 0.0021 |
| *CT → GR* | 0 – 0.05 | 0.0198 | 0.0003 | 0.0046 |
| *AE → CT* | 0 – 0.05 | 0.0128 | 0.0074 | 0.0045 |
| *CT → AE* | 0 – 0.05 | 0.001 | 0.0022 | 0.0046 |

**Supplementary Table 5** – System “Crete”; Point estimates and 95% confidence intervals from non-parametric bootstrapping for the best-fitting demographic scenario for system Crete – scenario C. Haploids/ generation in migration rates were estimated by multiplying each migration rate with the point estimate haploid size of the population of origin.

| Parameter | 95% Lower limit | Point estimate | 95% Upper limit |
| --- | --- | --- | --- |
| Haploid population sizes | |  |  |
| *GR* | 509 | 1’465 | 7’880 |
| *AE* | 383 | 802 | 60’962 |
| *CT* | 107 | 373 | 944 |
| Bottleneck intensity (haploid size) | | | |
| *AE* | 13 | 48 | 2922 |
| *CT* | 6 | 74 | 243 |
| Time of divergence (generations ago) | | | |
| *T^CRETE^* | 68 | 321 | 1’400 |
| Ancient Migration rates (forward migration in haploids/generation) | | | |
| *GR → AE* | 0.1421 | 1.3185 | 61.823 |
| *AE → GR* | 0.0441 | 0.0577 | 36.09 |
| Current Migration rates (forward migration in haploids/generation) | | | |
| *GR → AE* | 0.082 | 61.09 | 57.867 |
| *AE → GR* | 7.218 | 0.481 | 56.942 |
| *GR → CT* | 0.063 | 6.739 | 10.987 |
| *CT → GR* | 0.0268 | 0.7833 | 6.0426 |
| *AE → CT* | 0.1604 | 3.6892 | 8.3408 |
| *CT → AE* | 0.0373 | 1.6785 | 9.698 |

**Supplementary Table 6** – Parameter ranges and estimates of fastsimcoal2 for system “Cyprus”. Initial range is the range from which the software draws initial the parameter values. The distributions are uniform for every parameter except *Bottleneck Intensity* which is log-uniform. Highest likelihood estimates are provided for every parameter. In scenarios A, C the GH population diverges first from IS and therefore time of divergence for CY is scaled to be smaller and ancient migration is only between IS and GH. For scenario B, CY is more ancient than GH and the time of split is reversed, and ancient migration rates are between IS and CY. IS: Israel; GH: Ghost population; CY: Cyprus.

| Parameter | Initial range | Point estimates | | |
| --- | --- | --- | --- | --- |
|  |  | **Cyprus A** | **Cyprus B** | **Cyprus C** |
| Haploid population sizes | |  |  |  |
| *IS* | 5’000 – 150’000 | 5’140 | 7’959 | 5’631 |
| *GH* | 100 – 150’000 | 130’619 | 487 | 109’265 |
| *CY* | 100 – 50’000 | 122 | 47’095 | 292 |
| Bottleneck intensity (fraction of current population size) | | | |  |
| *GH* | 0.01 – 0.5 | 0.110 | 0.296 | 0.025 |
| *CY* | 0.01 – 0.5 | 0.104 | 0.135 | 0.023 |
| Time of divergence (proportion of 6000; generations) | | | |  |
| *T^CYPRUS^ (scenario A, C)* | 0.001 – 0.999 | 0.164 | - | 0.075 |
| *T^GHOST^ (scenario A, C)* | 0.001 – 3 | 1.709 | - | 0.715 |
| *T^CYPRUS^ (scenario B)* | 0.001 – 3 | - | 0.646 | - |
| *T^GHOST^ (scenario B)* | 0.001 – 0.999 | - | 0.198 | - |
| Ancestral migration rates (backwards in time) | | | | |
| *IS → GH (scenario A, C)* | 0 – 0.05 | 0.0001 | - | 0.0005 |
| *GH → IS (scenario A, C)* | 0 – 0.05 | 3e-05 | - | 0.0004 |
| *IS → CY (scenario B)* | 0 – 0.05 | - | 0.0431 | - |
| *CY → IS (scenario B)* | 0 – 0.05 | - | 0.0056 | - |
| Current migration rates (backwards in time) | | | | |
| *IS → GH* | 0 – 0.05 | 0.001 | 0.0001 | 2e-05 |
| *GH → IS* | 0 – 0.05 | 0.0087 | 0.0086 | 0.0039 |
| *IS → CY* | 0 – 0.05 | 0.0001 | 0.0001 | 0.0001 |
| *CY → IS* | 0 – 0.05 | 0.0078 | 0.0005 | 0.0152 |
| *GH → CY* | 0 – 0.05 | 0.0018 | 0.0094 | 0.0108 |
| *CY → GH* | 0 – 0.05 | 0.0424 | 0.035 | 0.0051 |

**Supplementary Table 7** – System “Cyprus”; Point estimates and 95% confidence intervals from non-parametric bootstrapping for the best-fitting demographic scenario for system Cyprus – scenario A. Haploids/ generation in migration rates were estimated by multiplying each migration rate with the point estimate haploid size of the population of origin.

| Parameter | 95% Lower limit | Point estimate | 95% Upper limit |
| --- | --- | --- | --- |
| Haploid population sizes | |  |  |
| *IS* | 5’068 | 5’140 | 19’159 |
| *GH* | 34’780 | 130’619 | 172’130 |
| *CY* | 101 | 122 | 785 |
| Bottleneck intensity (haploid size) | | | |
| *GH* | 1’176 | 14’423 | 11’739 |
| *CY* | 3 | 12 | 145 |
| Time of divergence (generations ago) | | | |
| *T^CYPRUS^* | 633 | 986.1 | 3’336 |
| *T^GHOST^* | 2’392 | 10’250.9 | 11’739 |
| Ancient Migration rates (forward migration in haploids/generation) | | | |
| *IS → GH* | 4.8948 | 0.97146 | 160.38 |
| *GH → IS* | 231 | 13.1 | 6440 |
| Current Migration rates (forward migration in haploids/generation) | | | |
| *IS → GH* | 5.11 | 44.944 | 224 |
| *GH → IS* | 0.2069 | 133.022 | 13.4799 |
| *IS → CY* | 27.5 | 40.16 | 171 |
| *CY → IS* | 0.00026 | 0.0122 | 0.02353 |
| *GH → CY* | 145.9 | 5538.598 | 51’880 |
| *CY → GH* | 0.113 | 0.216 | 42.9 |

**Supplementary Figures**

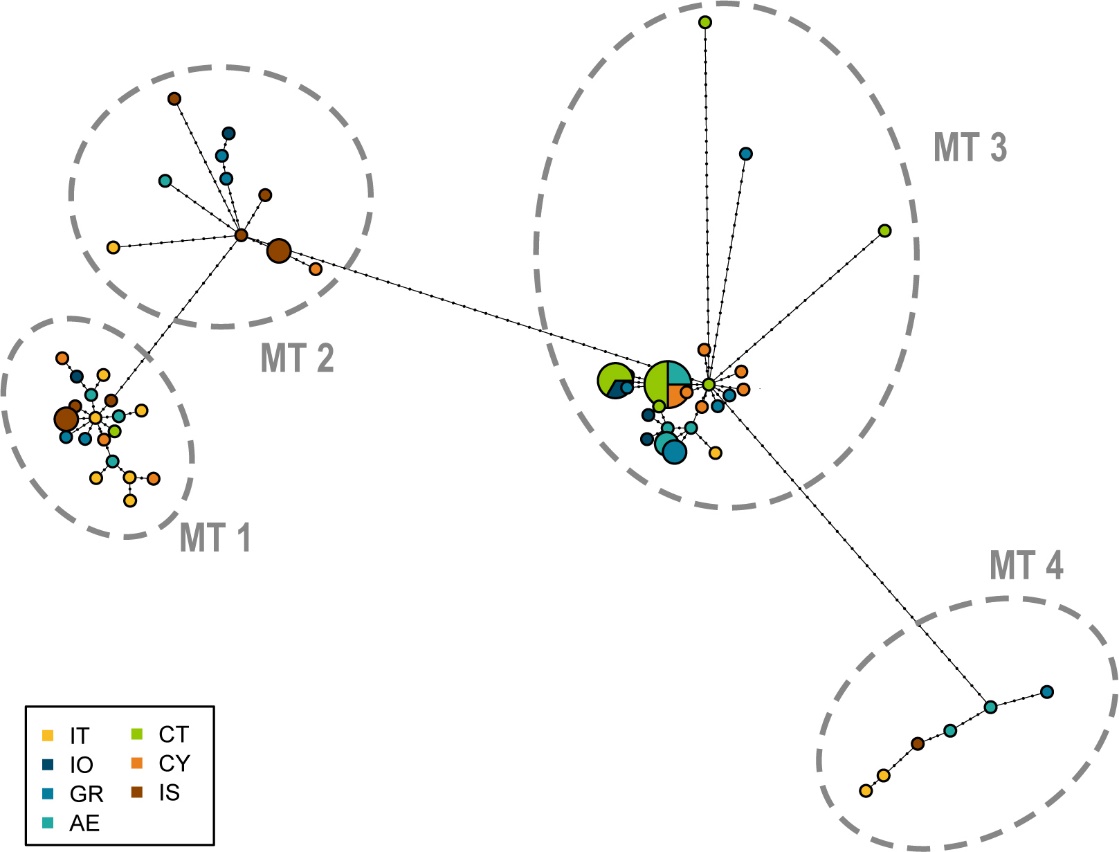

**Sup. Fig. 1 –** Haplotype network based on mtDNA of barn owls in the Mediterranean. The whole mitochondrial genome, except the D-loop, was used to construct the network. Small black dots represent individual mutation steps. Pie size is proportional to number of individuals carrying the haplotype. Large dashed circles indicate MT haplogroups 1 to 4, which are plotted in Figure 1a.

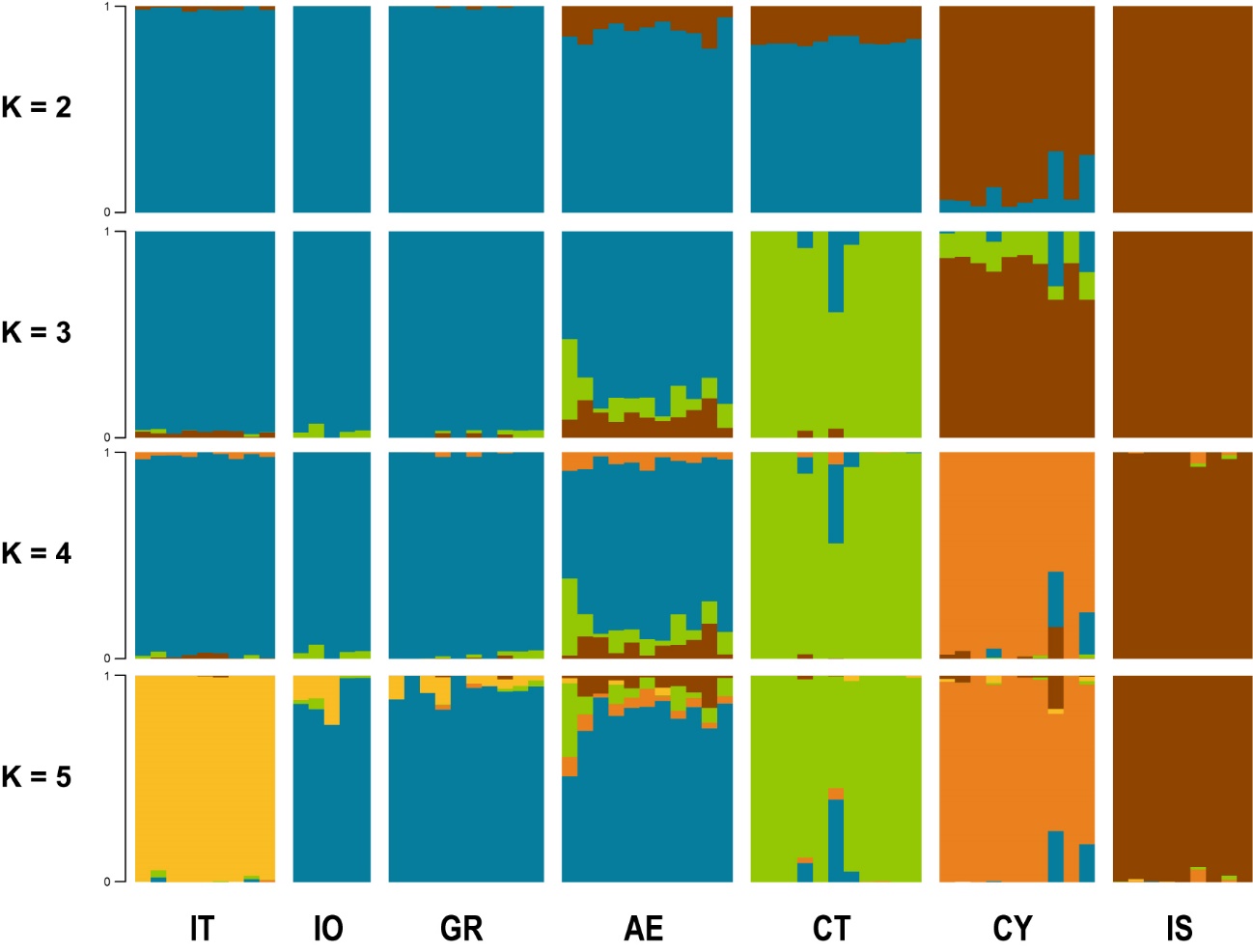

**Sup. Fig. 2 –** Individual clustering estimated by sNMF for K 2 to 5 lineages. Each vertical bar represents one individual, and the colours represent the relative contributions of each genetic lineage.

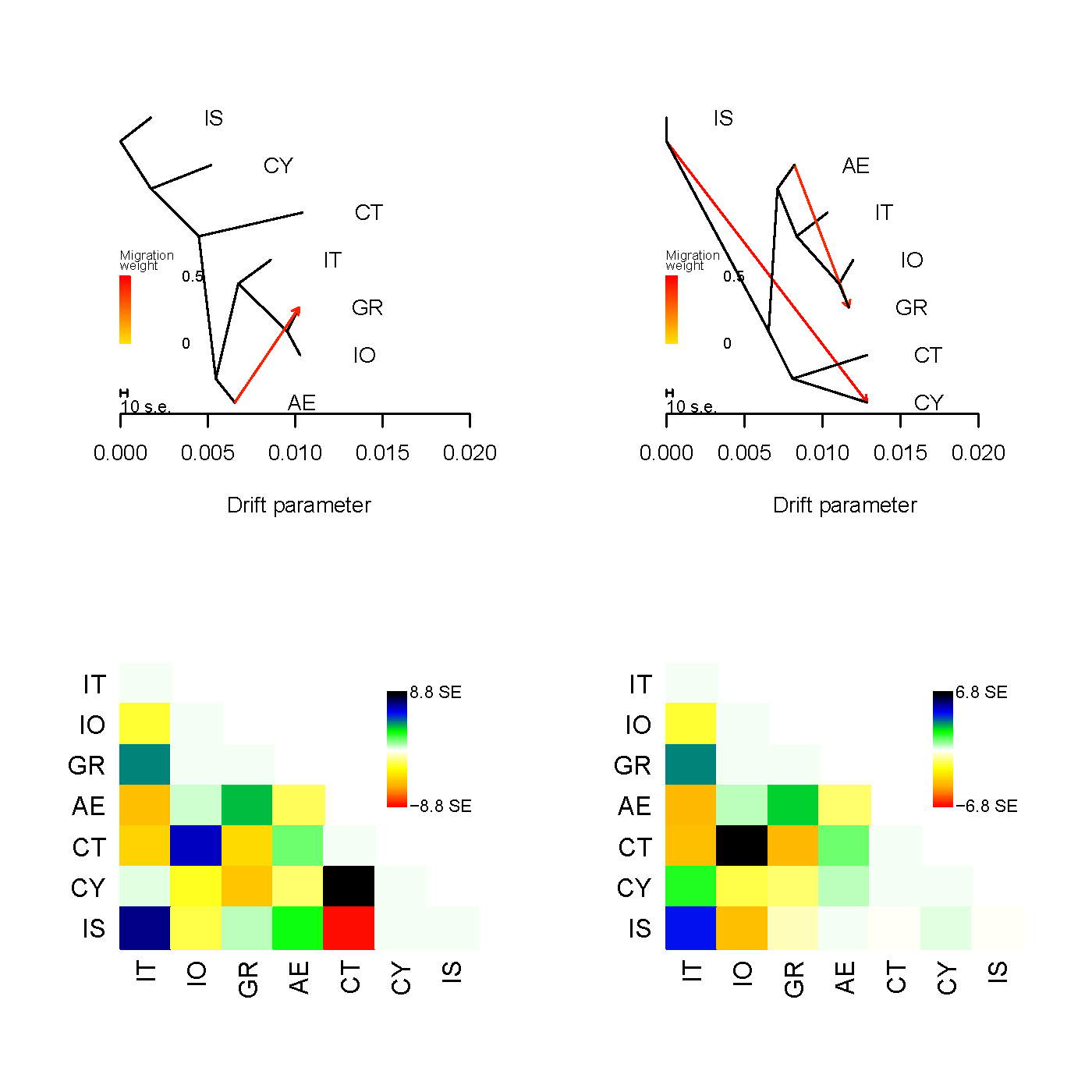

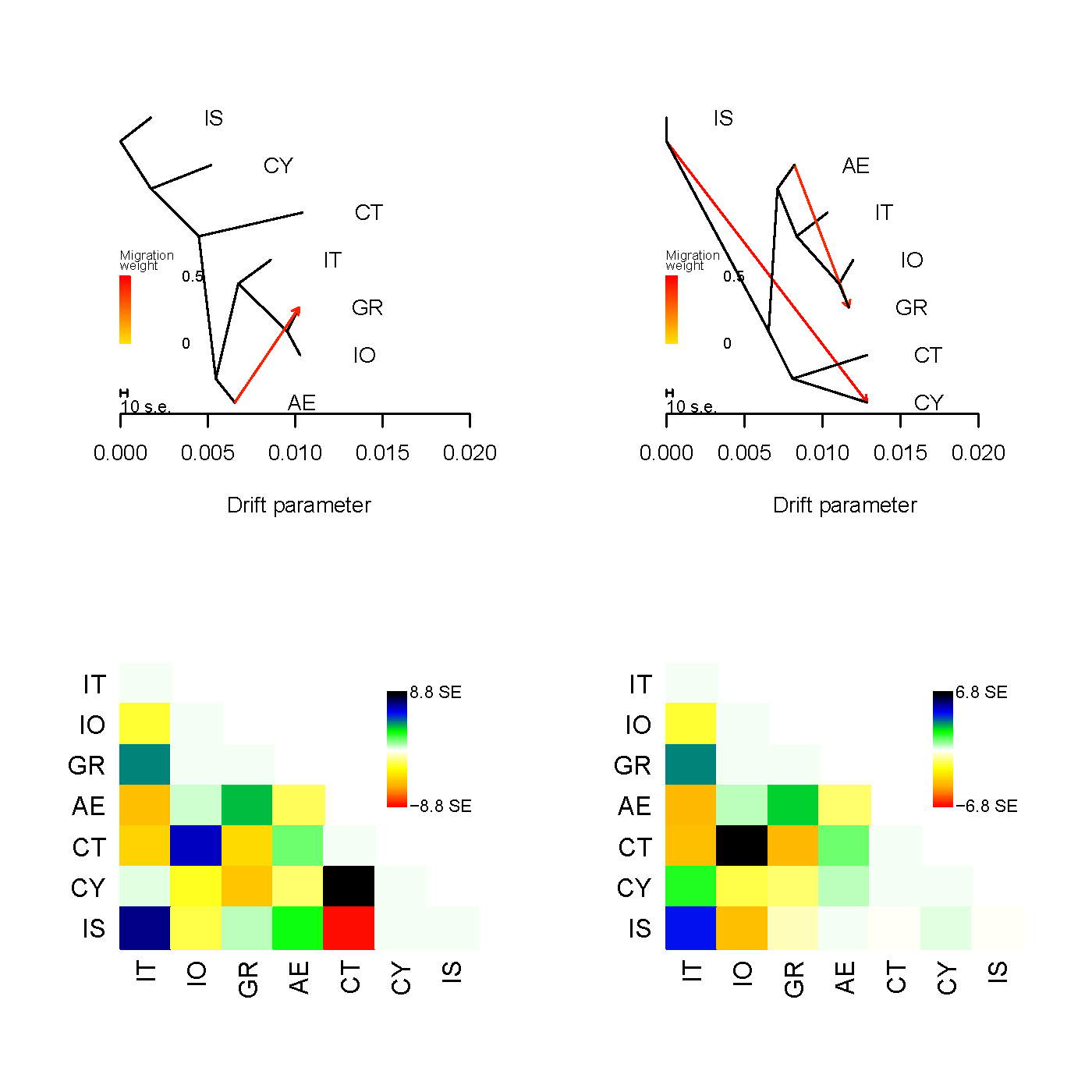

**Supplementary Figure 3** – Treemix analysis for 1 migration events with its residual matrix.

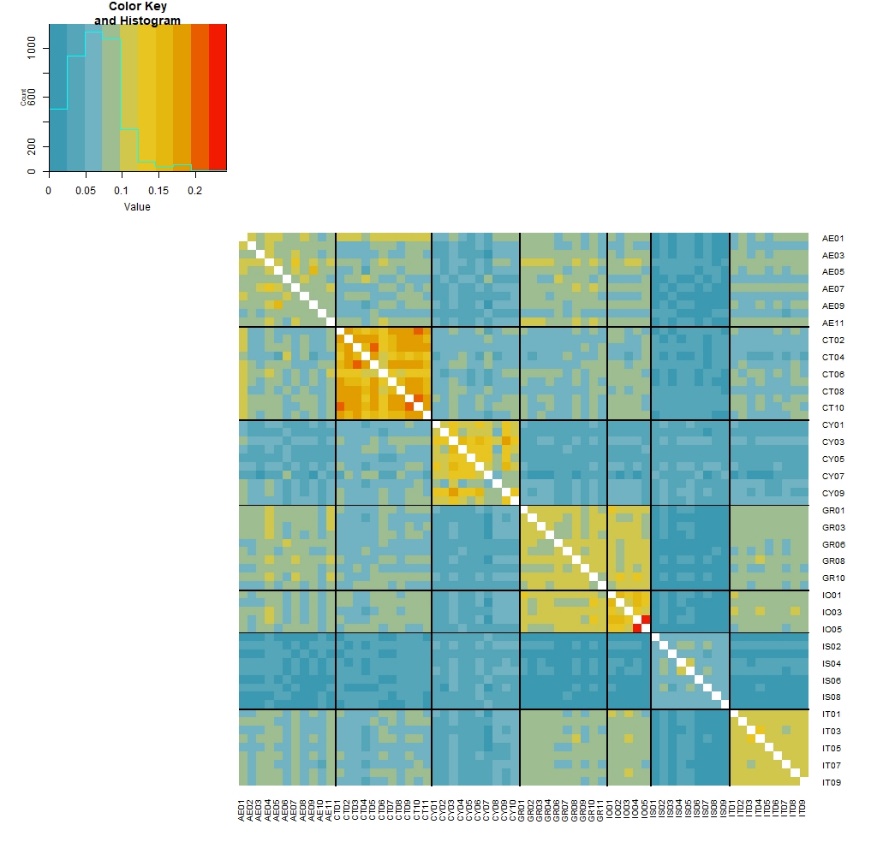

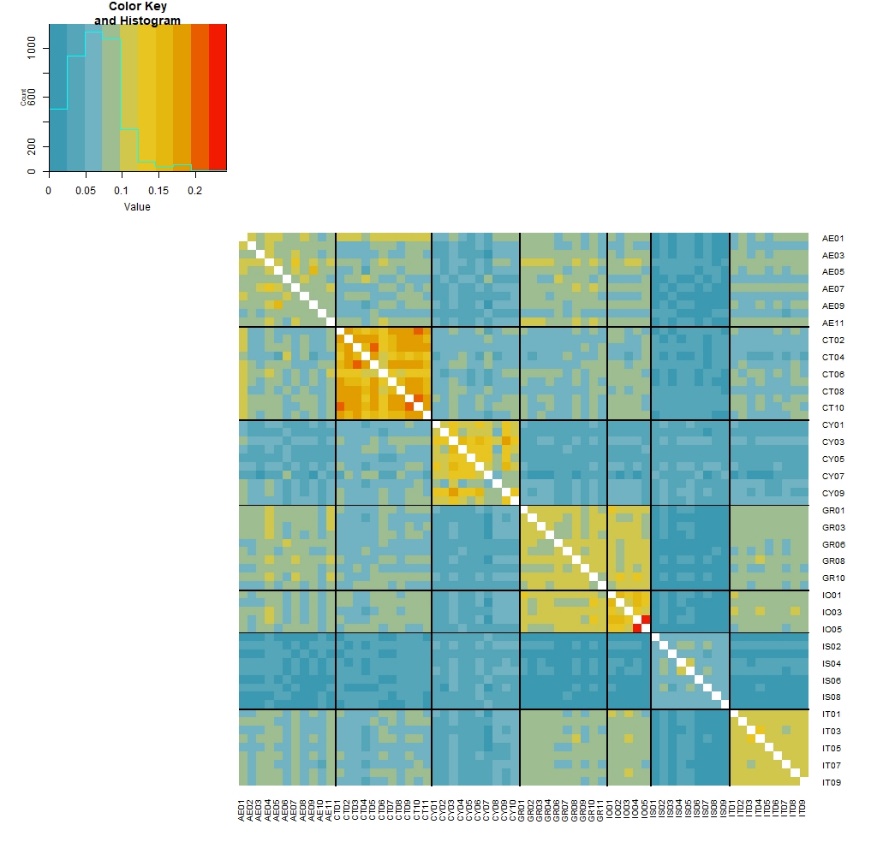

**Supplementary Figure 4** – Pairwise individual relatedness (β) heatmap between all individuals.

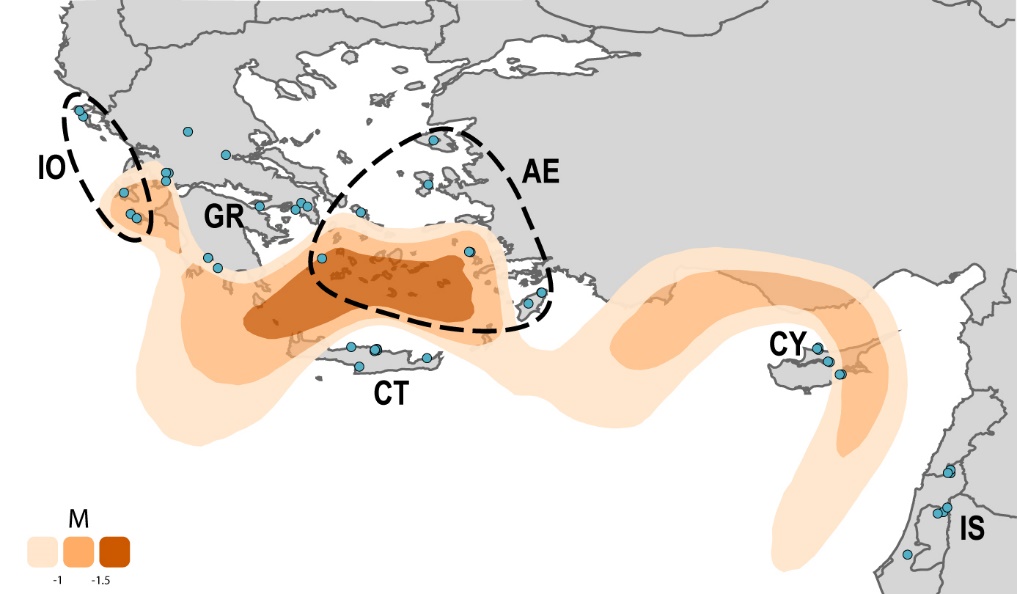

**Supplementary Figure 5** – Estimated effective migration surface (EEMS) based on whole-genome data. Orange shading denote regions of lower than average gene flow, darker shading indicates a stronger barrier. Dots indicate individual sampling location. Dashed lines delimit the Greek archipelagos sampled.

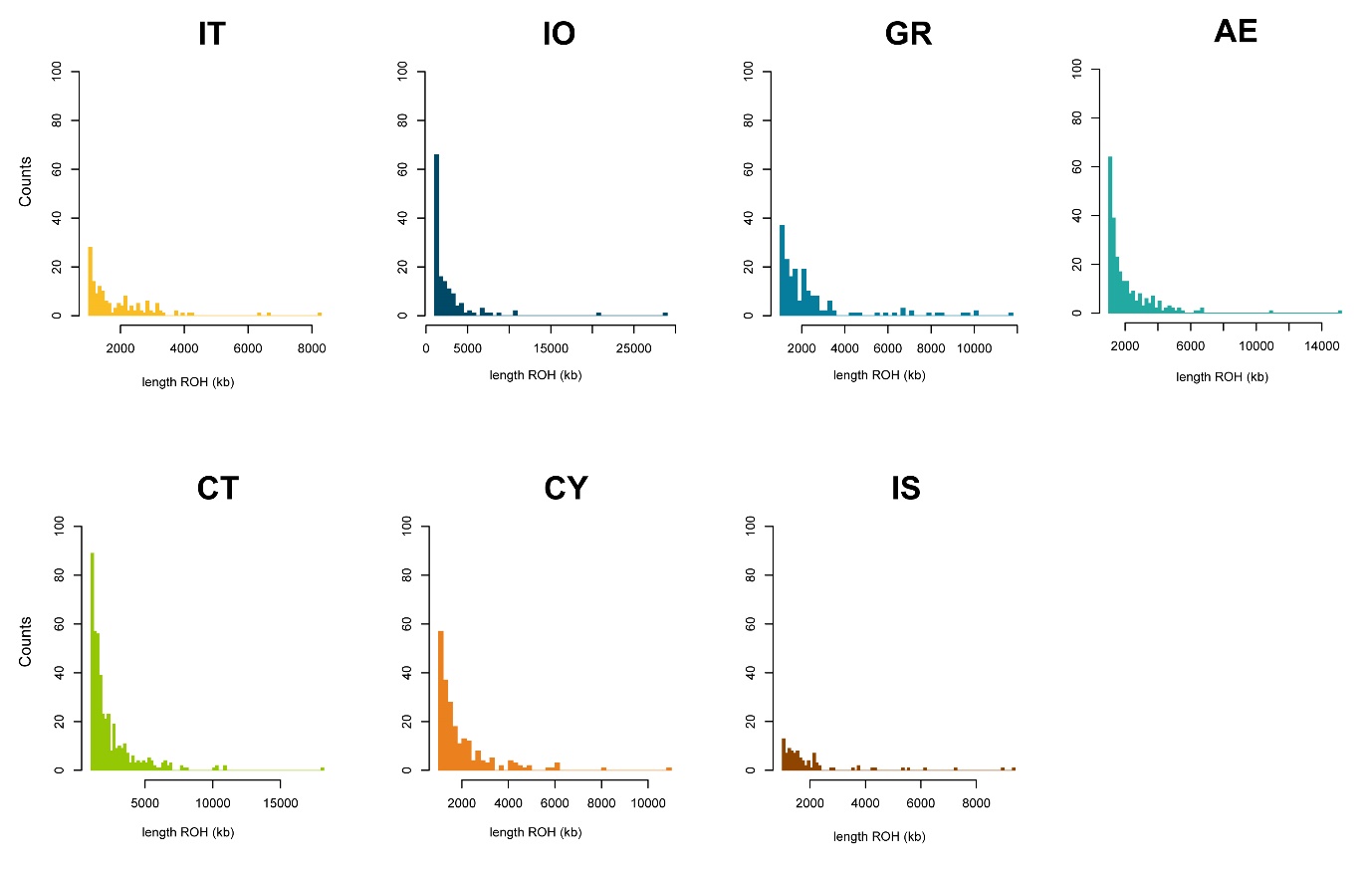

**Supplementary Figure 6** – Distribution of ROH segments per population. Vertical axes are the same between plots but note the differences in the horizontal axes.

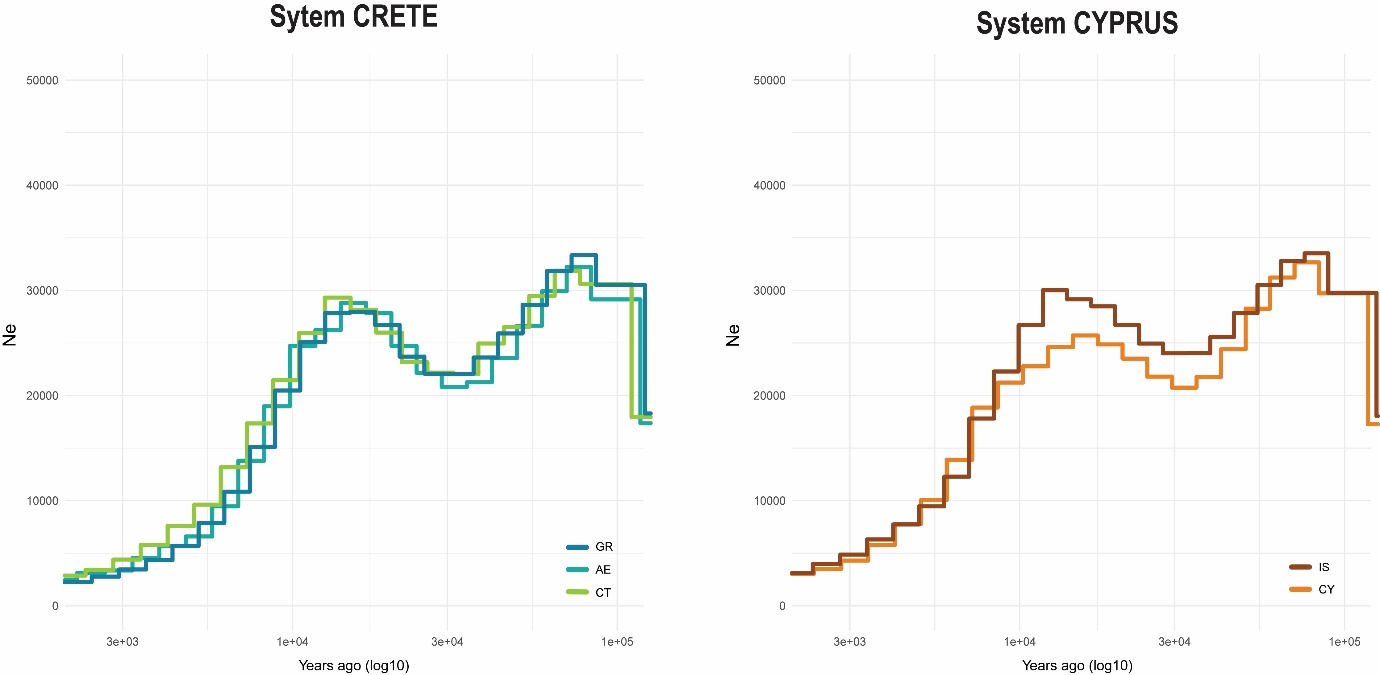

**Supplementary Figure 7** – PSMC results for system Crete and Cyprus, separated for readability. Each line represents the median across individuals in each population. Plotted with 8.28x10^-9^ mutation rate and 3.6 years generation time.
